# Transcriptome of the murine duodenum during recovery after a lethal dose of total body irradiation

**DOI:** 10.1101/2024.11.26.625550

**Authors:** Ling He, Kruttika Bhat, Frank Pajonk

## Abstract

In the aftermath of 9/11, the radiobiology community sought novel radiation mitigators capable of preventing death when administered 24 hours or later after exposure to lethal ionizing radiation. The survival and expansion of normal stem cells are crucial for restoring tissue integrity in time to prevent mortality. While FDA-approved drugs for acute radiation syndrome primarily target the hematopoietic system, restoring the integrity of the intestinal lining is equally important for survival. However, the radiation response of the intestinal stem cell (ISC) population and its niche environment is not as well understood as that of the bone marrow. The aim of this study was to explore early transcriptomic changes in the small intestine after a lethal dose of total body irradiation (TBI), and during subsequent recovery.

C3H/Sed/Kam mice were irradiated with a TBI dose of 16 Gy, the published LD_70/10_. The compound 1-[(4-Nitrophenyl)sulfonyl]-4-phenylpiperazine (NSPP) was administered 24 hours post-irradiation. RNAs from the proximal duodenum were extracted at 28, 72, and 96 hours post-irradiation and subjected to RNA-sequencing. Differentially expressed genes were analyzed using gene-set enrichment analysis.

Radiation induced significant transcriptomic changes known to precede the death of lymphatic endothelial and epithelial cells. Upregulation of *Lgr5+* ISC gene signature was observed during recovery. NSPP treatment further amplified the activation of ISC-associated genes and other regenerative markers. Notably, gene *Psrc1* showed strong activation throughout the recovery process, highlighting its potential role in this regenerative response. These findings suggest additional points of intervention for radiation mitigation in the intestines beyond targeting programmed cell death.

## Introduction

Exposure to ionizing radiation during reactor accidents, nuclear weapon detonations, or dirty bombs leads to a symptom complex that is summarized as acute radiation syndrome or ARS. Among the organ systems that determine the survival of the individual, the response of the small intestines has been well studied [1, 2]. The small intestines are organized hierarchically with *Lgr5+* crypt base columnar (CBC) cells wedged between Paneth cells at the base of the crypts forming the intestinal stem cell (ISC) population [3]. Progeny of CBC cells migrate in a conveyor-like fashion to the apex of the intestinal villi, where the cells are eventually shed off and die. Following ionizing radiation exposure, epithelial cell death in the small intestines occurs rapidly, leading to the loss of *villi* and failure of the intestine’s resorption and barrier functions [2]. This results in fluid loss, bleeding, and bacterial translocation, which can cause sepsis and death unless the surviving intestinal stem cells can reconstitute tissue integrity in time.

The ability of surviving CBC cells to repair intestinal tissue integrity is influenced not simply by the radiation dose received by the epithelial cells but also by the response of various other tissue components of the intestinal wall. For instance, the death of endothelial cells in the *villi* capillaries affects the radiation response of the small intestine [4], and crypts in proximity to Peyer’s patches are more radioresistant than those further from these immune hubs [5, 6]. However, the overall interplay of these multiple tissue components in response to radiation is incompletely understood.

In the event of mass casualties resulting from nuclear incidents, there is an urgent need for drugs capable of mitigating the effects of lethal radiation doses [7]. The logistical challenges, such as mass panic or infrastructure collapse, necessitate the development of drugs that can be administered up to 24 hours after exposure to a lethal dose of radiation (e.g., an LD_70/10_). Our previous efforts to identify such drugs have involved high-throughput screening of drug libraries, using lymphocyte death as the primary endpoint. Among the compounds identified, 1-[(4-Nitrophenyl)sulfonyl]-4-phenylpiperazine (NSPP) has shown the most efficacy in mitigating hematopoietic and gastrointestinal acute radiation syndrome (GI-ARS) [8]. However, this approach did not fully capture the complex responses of intestinal tissue to radiation exposure [9].

In this study, we performed a comprehensive gene expression analysis of the murine duodenum in C3Hf/Sed/Kam mice to uncover the early transcriptomic changes triggered by a lethal dose of total body irradiation (TBI) and the subsequent recovery processes. Our analysis revealed dynamic shifts in gene expression over time, with early activation of pathways associated with cell death, cell cycle arrest, compromised hematopoiesis, and immune deficiencies. These early disruptions were then gradually subsided/restored, followed by the upregulation of genes involved in cell-cell adhesion, protein digestion and absorption, critical signaling pathways, and ISC-associated gene signatures.

Importantly, treatment with NSPP further enhanced the activation of ISC signatures, expediting tissue repair and recovery. Notably, the expression of *Psrc1* was significantly elevated with NSPP treatment, suggesting its pivotal role in promoting the regenerative effects observed. These findings highlight the complex and coordinated response of intestinal tissues to radiation-induced damage and underscore the therapeutic potential of NSPP in mitigating GI-ARS. By advancing our understanding of the molecular mechanisms involved in intestinal recovery, this study lays the groundwork for developing more effective and innovative radiation countermeasures.

## Material and Methods

### Animals

C3Hf/Sed//Kam mice were bred and maintained in a pathogen-free environment in the American Association of Laboratory Animal CARE-accredited Animal Facilities of Department of Radiation Oncology, University of California (Los Angeles, CA) in accordance with all local and national guidelines for the care of animals and was approved by the University of California Los Angeles’ Institutional Animal Care and Use Committee (IACUC).

### Irradiation

Twelve-week-old female C3Hf/Sed/Kam mice were irradiated using an experimental X-ray irradiator (Gulmay Medical Inc. Atlanta, GA) at a dose rate of 2.8159 Gy/min for the time required to apply a prescribed dose. The X-ray beam was operated at 300 kV and hardened using a 4 mm Be, a 3 mm Al, and a 1.5 mm Cu filter. NIST-traceable dosimetry was performed using a micro-ionization chamber and dosimetry was confirmed using films (Ashland Specialty Ingredients G. P., Bridgewater, NJ) placed at 5 positions inside the abdomen of mouse cadavers.

### In vivo drug administration

Twenty-four hours after irradiation, the mice received three daily subcutaneous injections of 5 mg/kg 1-[(4-Nitrophenyl)sulfonyl]-4-phenylpiperazine (NSPP, Vitascreen, Champaign, IL). This compound was dissolved in 15 µl DMSO and then suspended in 1 ml of 1% Cremophor EL, with each injection spaced 24 hours apart. Control animals received injections of the DMSO/Cremophor EL solution without the compound.

### RNA isolation

Twenty-eight, seventy-two and ninety-six hours after irradiation or sham irradiation the animals were euthanized using isoflurane and cervical dislocation. The abdominal cavity was immediately opened, and small intestines mobilized, harvested and flushed with PBS. Even under homeostasis the activity of signaling pathways changes dramatically along the murine small intestines [10]. In order to minimize this effect, we restricted the analysis to the duodenum. The proximal half of the duodenum of each mouse was cut, minced, and RNA was extracted using Trizol.

### Bulk RNA sequencing

Bulk RNA sequencing (RNASeq) was performed by Novogene (Chula Vista, CA). Quality and integrity of total RNA was controlled on Agilent Technologies 2100 Bioanalyzer (Agilent Technologies; Waldbronn, Germany). The RNA sequencing library was generated using NEBNext® Ultra RNA Library Prep Kit (New England Biolabs) according to manufacturer’s protocols. The library concentration was quantified using a Qubit 3.0 fluorometer (Life Technologies), and then diluted to 1 ng/uL before checking insert size on an Agilent Technologies 2100 Bioanalyzer (Agilent Technologies; Waldbronn, Germany) and quantifying to greater accuracy by quantitative Q-PCR (library molarity >2 nM). The library was sequenced on Illumina NovaSeq6000 with an average of 20M reads per RNA sample. Reads were then mapped to the mouse genome (mg39) following their standard pipeline [11].

Read counts were analyzed using the iDEP package (version 2.0) [12]. Differentially expressed genes were calculated using the DESeq2 algorithm with a minimum of a 2-fold change and a false discovery rate (FDR) of 0.1. Enrichment *p*-values were calculated based on a one-sided hypergeometric test. *P*-values were then adjusted for multiple testing using the Benjamini-Hochberg procedure and converted to FDR. Fold Enrichment was defined as the percentage of genes in the list belonging to a pathway, divided by the corresponding percentage in the background.

### TdT In Situ Apoptosis Detection

Twenty-four hours after total body irradiation with 16 Gy, the mice were euthanized, and their small intestines were immediately harvested and fixed in formalin. The tissues were then embedded in paraffin, and 4 µm sections were prepared for TdT In Situ Apoptosis Detection (#4812-30-K, R&D Systems, Minneapolis, MN). Briefly, the slides were baked for one hour in an oven at 65 °C, then dewaxed in two successive xylene baths for 5 minutes each, followed by hydration through an alcohol gradient (ethanol 100%, 90%, 70%, 50%, 25%) for 5 minutes each. The sections were subsequently processed for labeling and viewing as per the manufacturer’s instructions.

### Statistics

All data presented are representative of at least three biologically independent experiments. For bulk RNA sequencing, differentially expressed genes were identified with a minimum fold-change threshold of 2 and a false discovery rate cut-off of 0.1.

## Results

### Apoptosis induction in small intestines by radiation

In this study we irradiated C3Hf/Sed/Kam mice with the published LD_70/10_ of 16 Gy for total body irradiation [13]. This dose results in the manifestation of a full GI-ARS, with 70% of the mice expected to die within 10 days post-irradiation. In mice succumbing to GI-ARS the radiation-induced damage is so extensive that the subsequent tissue repair mechanisms in the small intestines are inadequate for recovery **(Figure 1a/b)**. We initially assessed the status of apoptosis induction in these mice 24 hours after irradiation, as this marks the start of radiation mitigator treatment. TUNEL staining revealed an extensive induction of apoptosis in the epithelial layer of the intestines including the stem cell-harboring crypts **(Figure 1c/d)**. Yet, our previous studies provided proof-of-concept that radiation mitigation of GI-ARS at 24 hours after radiation exposure is feasible [5, 8]. This suggested that radiation injury following a lethal dose of radiation initiates a tissue repair program in the small intestines that is insufficient for recovery but can be enhanced by radiation mitigators to improve survival.

**Figure 1.**
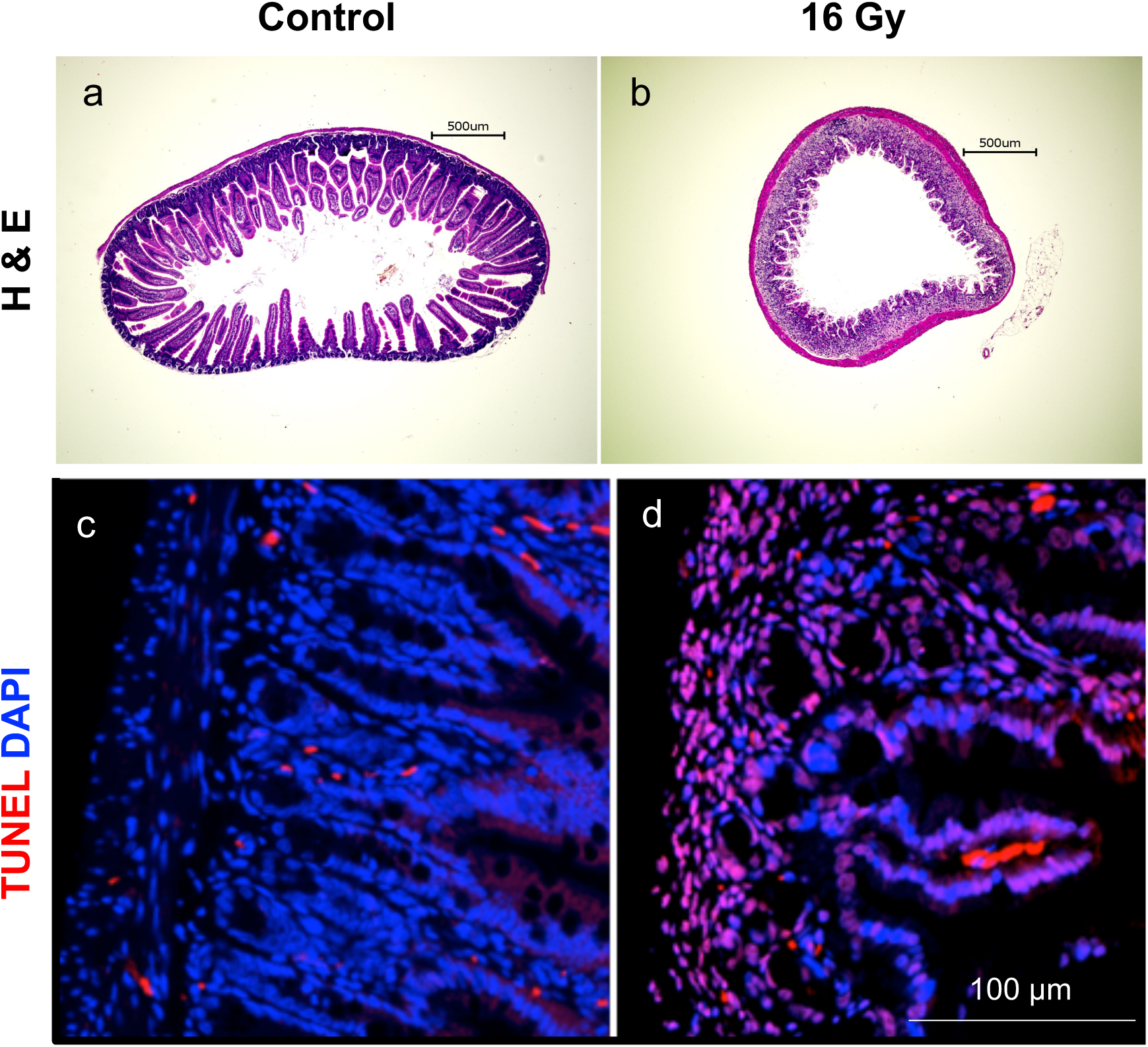
Apoptosis induction in murine small intestines by radiation. Total body irradiation of C3Hf/Sed//Kam mice with 16 Gy leads to the breakdown of the epithelial layer **(b)** and a massive induction of TUNEL-positive cells **(d)** at 24 hours post-irradiation, compared to sham-irradiated controls **(a/c)**.

### Total body irradiation induces distinct gene expression profiles in murine small intestines

To better understand the effects of radiation mitigators on the recovery process, we first examined early transcriptomic changes in the small intestines following a lethal dose of TBI, as well as changes during subsequent recovery without interventions. We performed bulk RNA sequencing at three time points (28, 72, and 96 hours post-TBI), as shown in **Figure 2a**. Principal component analysis demonstrated that biologically independent replicates clustered closely, indicating small variation between individual animals **(Figure 2b)**. Hierarchical clustering of differentially expressed genes (DEGs) revealed distinct gene expression profiles as early as 28 hours post-TBI **(Figure 2c)**, with 178 genes upregulated and 568 genes downregulated compared to the sham-irradiated controls **(Figure 2d)**. Next, we conducted a KEGG (Kyoto Encyclopedia of Genes and Genomes) enrichment analysis of the DEGs. The analysis revealed that the up-regulated genes significantly overlapped with gene sets involved in the *P53* signaling pathway and cell cycle regulation **(Figure 2e)**. This finding aligns with the established understanding of the biological response to DNA damage and stress caused by radiation, which includes the activation of *P53* signaling to mediate cell cycle arrest, DNA repair, or apoptosis [14]. However, the down-regulated genes were significantly associated with gene sets involved in hematopoietic cell lineage, primary immunodeficiency, and T cell receptor signaling pathway **(Figure 2f)**. This observation was in line with the well-known impact of radiation on the immune system, as it often results in compromised hematopoiesis and immune deficiencies [15] and the role of the intestines as a major part of the immune system.

**Figure 2.**
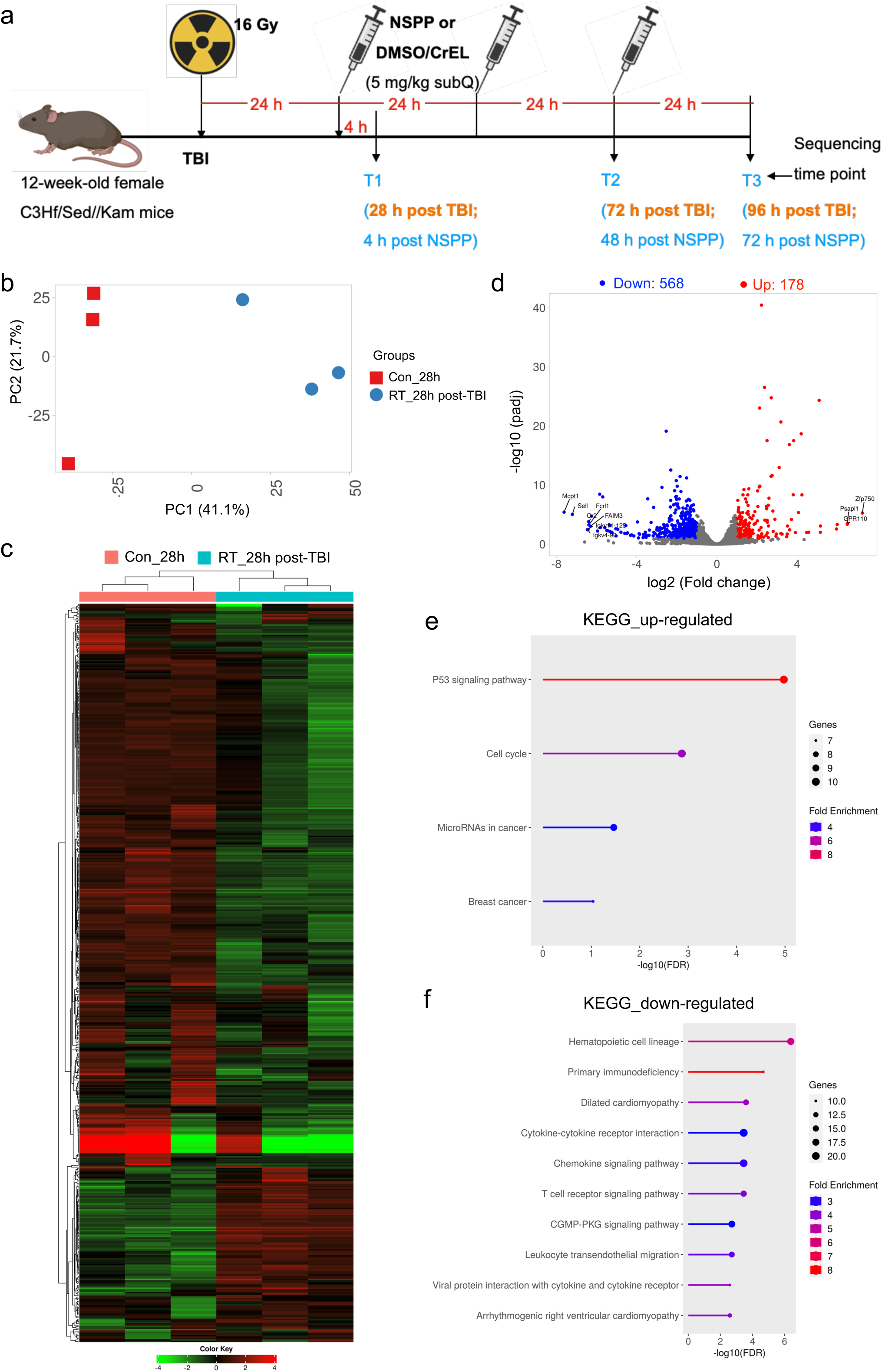
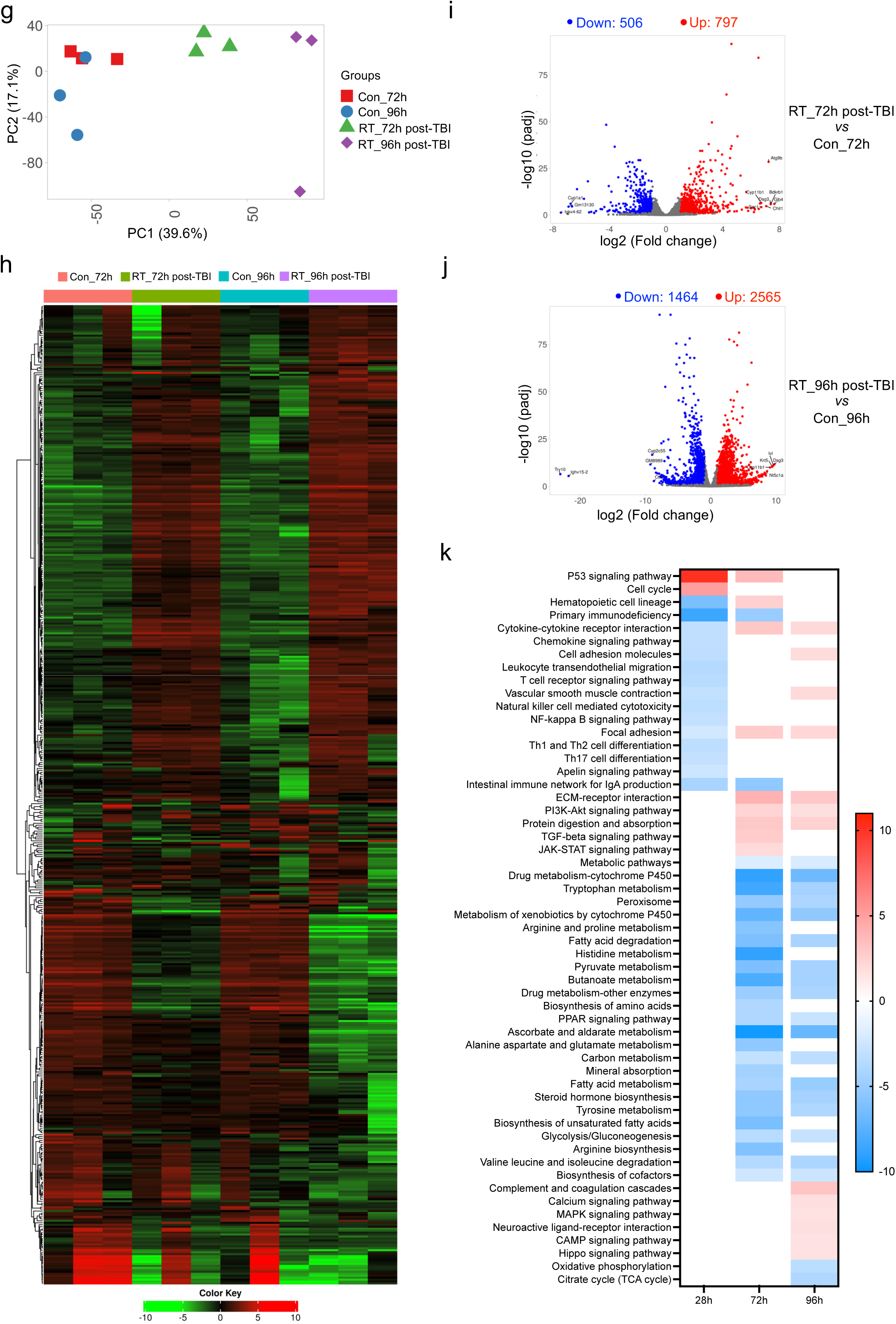
Total body irradiation induces distinct gene expression profiles in murine small intestines. Schematic representation of the experimental design **(a)**. Principal component analysis **(b)**, heatmap with hierarchical clustering of gene expression profiles **(c)**, and a volcano plot illustrating differentially expressed genes (DEGs) **(d)** in murine small intestines 28 hours post-TBI. KEGG gene set enrichment analysis of upregulated **(e)** and downregulated **(f)** differentially expressed genes at the same time point. Principal component analysis, heatmaps with hierarchical clustering, and volcano plots of DEGs at 72 and 96 hours post-TBI **(g–j)**. KEGG enrichment analysis of DEGs across all three time points compared to their corresponding sham-irradiated controls **(k)**.

To explore the subsequent transcriptomic changes during the recovery process, we analyzed samples at 72 and 96 hours post-irradiation. Again, principal component analysis showed that biologically independent replicates clustered closely, with distinct separation between different groups **(Figure 2g)**. Hierarchical clustering of DEGs revealed unique profiles among the groups **(Figure 2h)**. At 72 hours post-irradiation, 797 genes were upregulated and 506 genes were downregulated compared to the sham-irradiated control animals **(Figure 2i)**. At 96 hours post-irradiation 2565 genes were upregulated and 1464 genes were downregulated compared to the sham-irradiated controls **(Figure 2j)**. This indicated dynamic transcriptomic changes during the recovery phase.

To gain a better understanding of the dynamic transcriptomic changes, we conducted KEGG enrichment analysis of DEGs from all three time points post-irradiation compared to the corresponding sham-irradiated groups. Our findings showed that the initially upregulated *P53* signaling pathway and cell cycle modulation gradually subsided with completion of DNA repair. Concurrently, the downregulated pathways related to hematopoietic cell lineage and immunodeficiencies showed a gradual restoration over time, indicating recovery of immune function and the residing hematopoietic cell compartment **(Figure 2k)**. Additionally, we observed an upregulation in pathways related to cytokine-cytokine receptor interaction, cell adhesion molecules, focal adhesion, ECM-receptor interaction, and protein digestion and absorption during the recovery process, with a more pronounced increase at 96 hours post-irradiation **(Figure 2k)**. This aligns with the gradual recovery of the intestinal epithelial lining, which is crucial for nutrient absorption. The enhanced expression of these pathways suggests a robust effort by the tissues to restore integrity and function, facilitating overall recovery from radiation-induced damage. Conversely, we noted downregulation of metabolic pathways during the intestinal recovery process post-irradiation, indicating a prioritization of repairing damaged tissues and restoring cellular function over energy-consuming metabolic activities. Furthermore, we also observed activation of the PI3K-Akt, TGF-beta, JAK-STAT, MAPK, cAMP, and Hippo signaling pathways during the recovery process, with a more pronounced signature at 96 hours post-irradiation **(Figure 2k)**, indicating a coordinated activation of multiple signaling pathways known to support tissue repair and regeneration [16, 17].

### Upregulation of the Lgr5+ Intestinal Stem Cell Gene Signature During Recovery

The gene signature of *Lgr5^+^* ISCs has been extensively studied and characterized [18–20]. Integrating transcriptomics with proteomics analyses defined a molecular signature for *Lgr5^+^* CBC cells, consisting of 510 stem cell-enriched genes [18]. To extend our analysis, we compared our lists of DEG from three post-irradiation time points with these ISC signature genes. We found that several of these signature genes were upregulated during the recovery process, with the expression of an increasing number of genes becoming more pronounced over time **(Figure 3)**. This indicated active participation of *Lgr5^+^* ISCs in tissue regeneration and repair. Notably, the gene *Psrc1* consistently showed strong activation throughout the recovery process, highlighting its potential role in this regenerative response.

**Figure 3.**
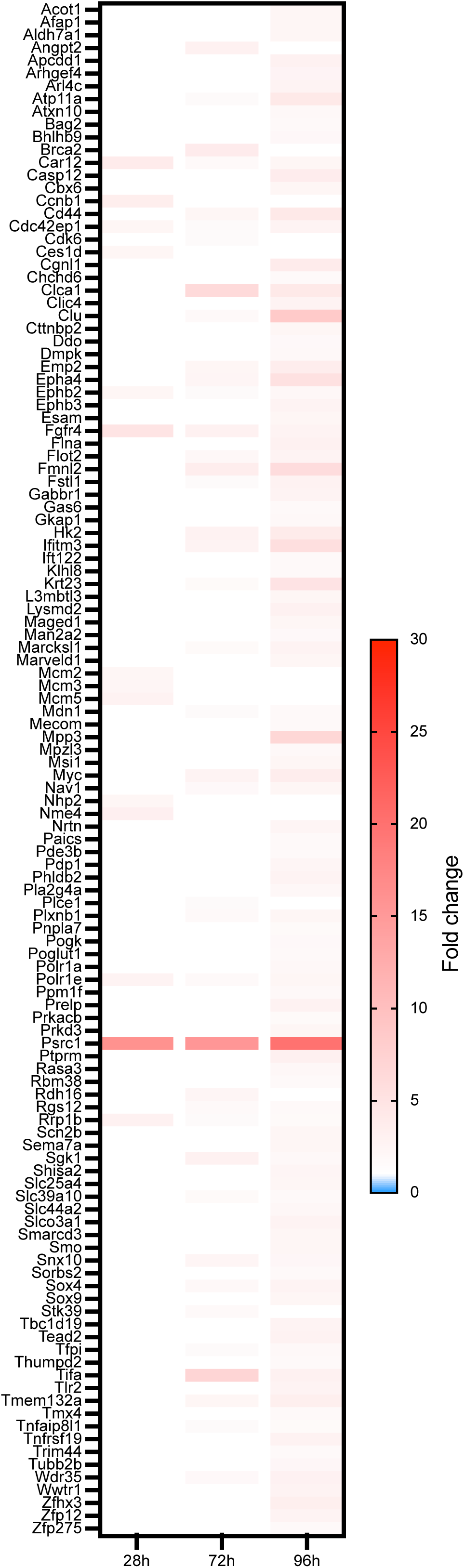
Upregulation of the *Lgr5+* Intestinal Stem Cell Gene Signature During Recovery. Heatmap displaying the enrichment of *Lgr5+* intestinal stem cell-associated genes among the up-regulated differentially expressed genes in murine small intestines at 28, 72, and 96 hours post-TBI.

Recently, two landmark studies published in *Cell* provided significant insights into the biology of intestinal stem cells [21, 22]. These studies identified and characterized the presence of multipotent intestinal stem cells located in the upper crypt zone. These undifferentiated cells play a pivotal role in maintaining intestinal homeostasis [21] and facilitating regeneration after irradiation-induced injury [22]. The cells, marked by *Fgfbp1*, have been shown to repopulate the intestinal epithelium, including regenerating the well-known *Lgr5^+^* crypt base columnar cells. Additionally, the sequencing data from the isthmus progenitor cells uncovered a novel signature of genes associated with intestinal stemness. To explore potential overlaps, we compared our DEG lists to their defined stemness signature. Interestingly, we found that two genes, *Areg* and *Clu*, were prominently present in our DEG lists at both 72-hour and 96-hour post-TBI time points, suggesting their potential involvement in the regenerative response following TBI.

*Early transcriptomic changes in the small intestines following NSPP mitigation* Previously, we have reported that NSPP has the potential to expand the intestinal stem cell pool, increase the number of regenerating crypts and prevent the GI-ARS [11]. To identify novel intervention targets influenced by NSPP that benefit the subsequent recovery process, we first explored the early transcriptomic changes at 28 hours post-TBI (4 hours after NSPP application). Again, principal component analysis showed close clustering of biologically independent replicates, with distinct separation between groups **(Figure 4a)**. Hierarchical clustering of DEGs revealed unique gene expression profiles among the groups **(Figure 4b)**. However, the drug effect at this early time point was minimal, with only 7 differentially downregulated genes in the NSPP plus radiation group compared to radiation alone **(Figure 4c)**, and 14 upregulated and 2 downregulated genes in the NSPP-only group compared to the sham-irradiated group **(Figure 4d)**. We also compared the NSPP-treated animals to the sham-irradiated group, identifying 222 differentially upregulated and 373 downregulated genes, as shown in the volcano plot **(Figure 4e)**. KEGG enrichment analysis of these DEGs revealed a significant overlap with gene sets involved in hematopoietic cell lineage and the intestinal immune network for IgA production **(Figure 4f)**. This suggests that NSPP may potentially boost intestinal recovery not only by increasing the number of ISCs but also by recovering resident immune cells, a function of NSPP described previously [8].

**Figure 4.**
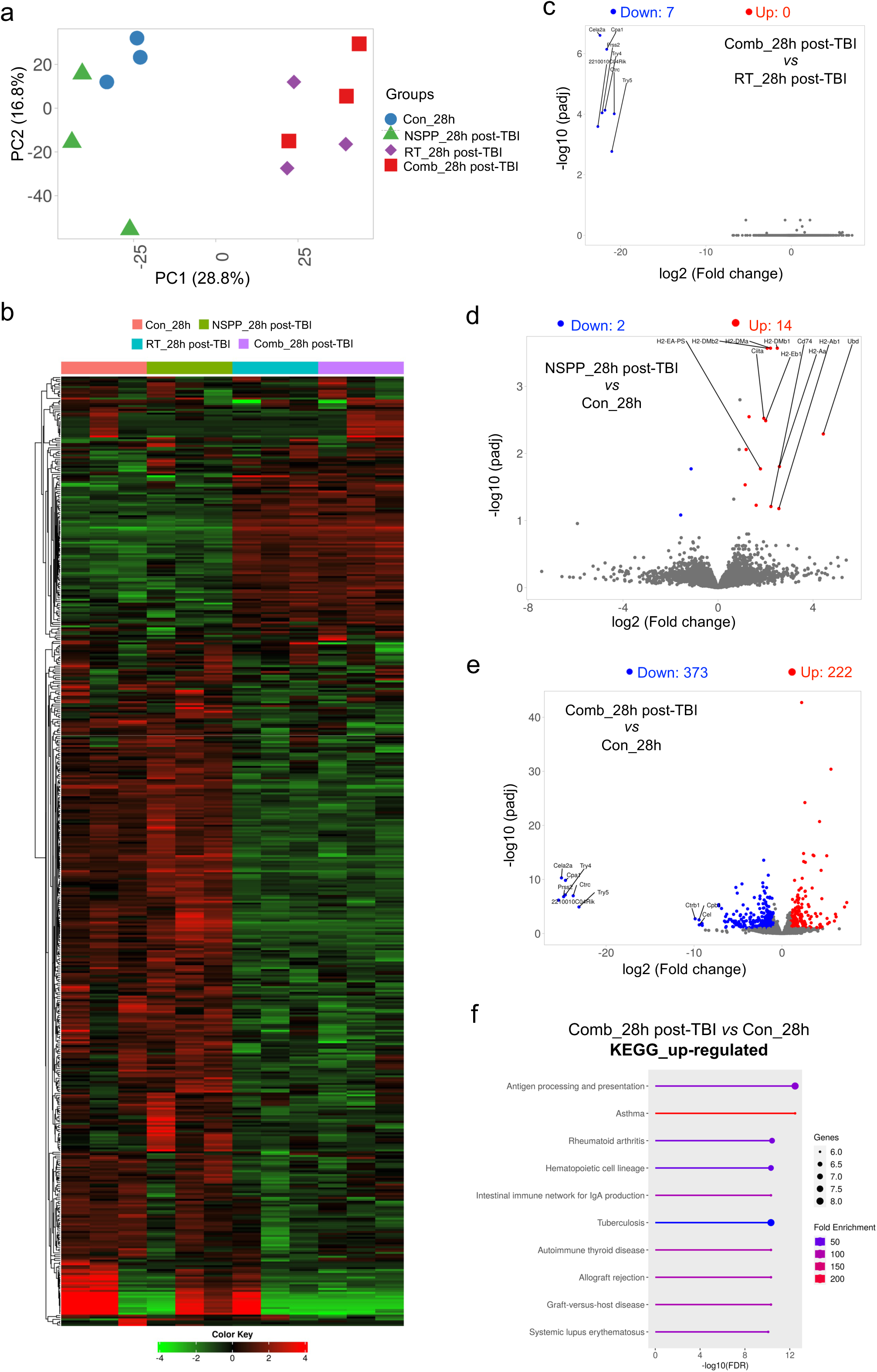
Early transcriptomic changes in the small intestines following NSPP mitigation. Principal component analysis **(a)**, and heatmap with hierarchical clustering of gene expression profiles **(b)** in murine small intestines at 28 hours post-TBI with or without NSPP treatment. Volcano plots displaying upregulated and downregulated DEGs across the treatment groups **(c-e)**. KEGG enrichment analysis of DEGs in the NSPP plus radiation group compared to sham-irradiated control at 28 hours post-TBI (4 hours after NSPP application) **(f)**.

### Transcriptomic changes during recovery following NSPP mitigation

Next, we examined transcriptomic changes at 72 hours (48 hours after NSPP application) and 96 hours (72 hours after NSPP application) post-TBI to identify potentially novel targets that facilitate recovery. Again, principal component analysis revealed distinct separation between groups **(Figure 5a),** and hierarchical clustering of DEGs revealed distinct gene expression profiles for the NSPP-treated animals **(Figure 5b)**. At 72 hours post-TBI, differences were subtle, with only 2 upregulated genes and 3 downregulated genes in the NSPP plus radiation group compared to radiation alone **(Figure 5c)**, and 11 downregulated genes in the NSPP-only group compared to sham-irradiated control **(Figure 5d)**. Comparing the NSPP plus radiation group to the sham-irradiated control, we found 922 upregulated and 517 downregulated genes **(Figure 5e)**. KEGG enrichment analysis revealed significant overlap of upregulated DEGs with gene sets involved in focal adhesion, cytokine-cytokine receptor interaction, ECM-receptor interaction, and PI3K-Akt signaling pathway **(Figure 5f/g)**. In contrast, downregulated genes overlapped with metabolism-related pathways, consistent with observations from the radiation-only group compared to the sham-irradiated group in **Figure 2k**.

**Figure 5.**
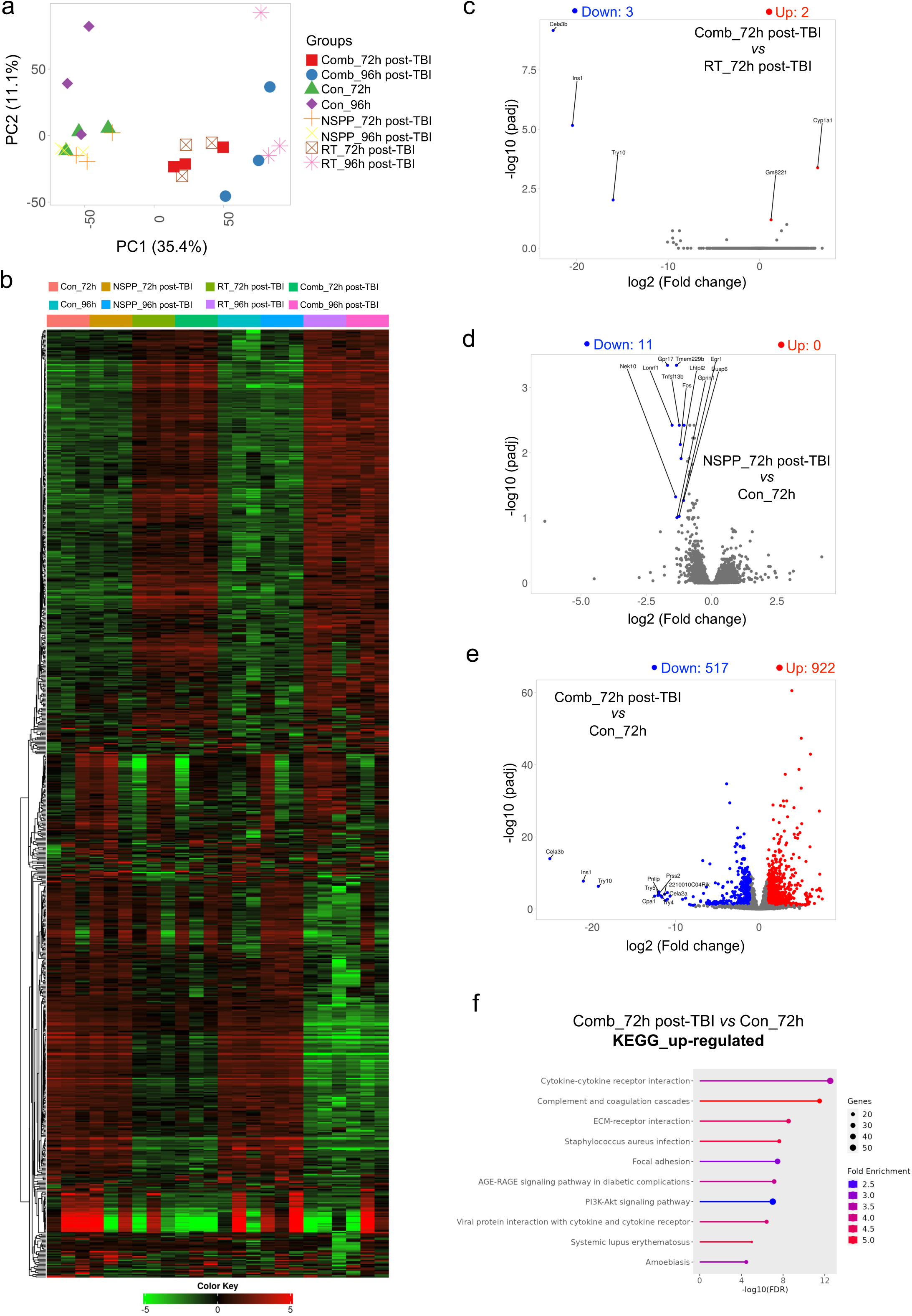
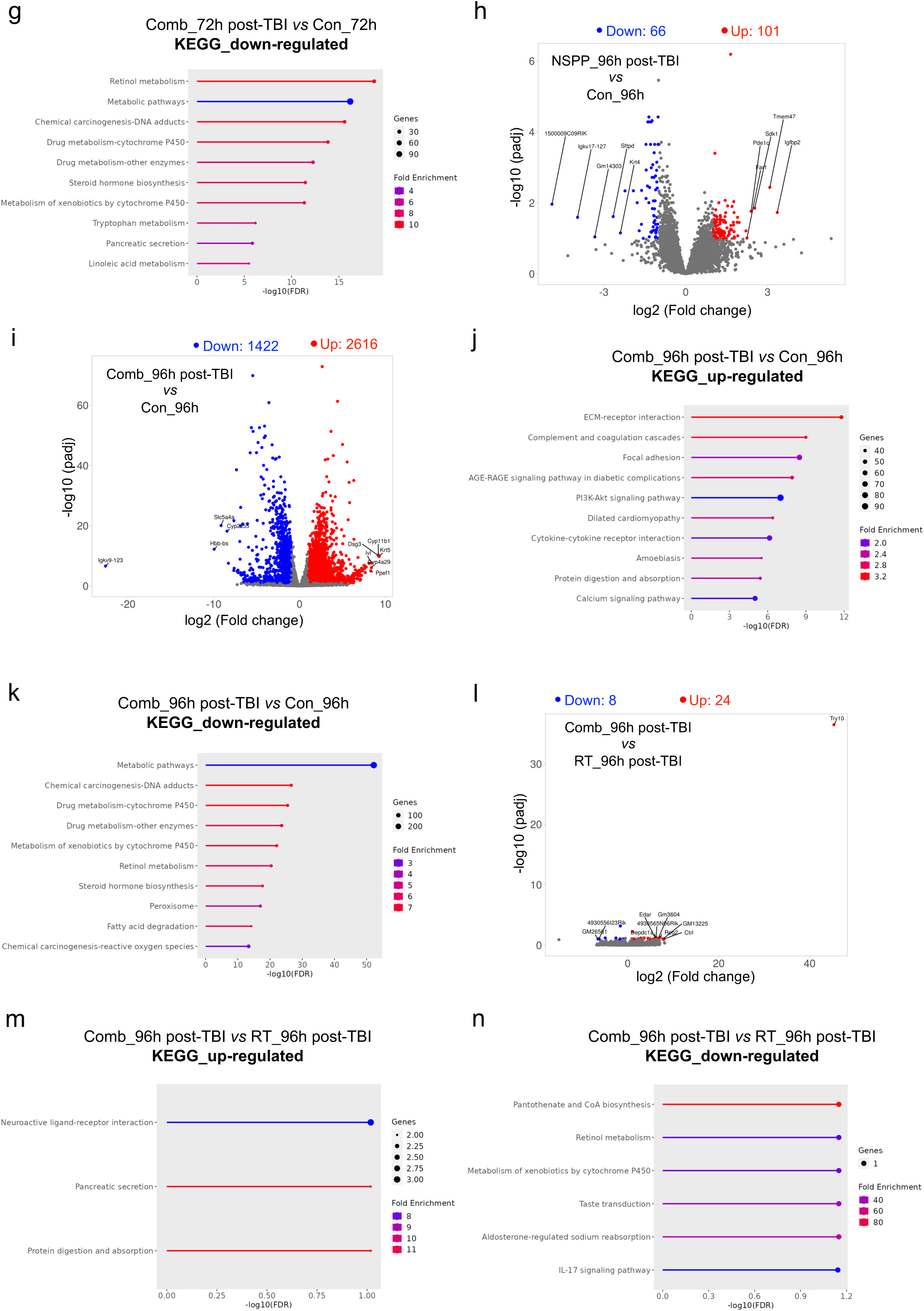
Transcriptomic changes during recovery following NSPP mitigation. Principal component analysis **(a)**, and heatmap with hierarchical clustering of gene expression profiles **(b)** in murine small intestines at 72 and 96 hours post-TBI with or without NSPP mitigation. Volcano plots showing upregulated and downregulated DEGs across all treatment groups at 72 hours post-TBI **(c-e)**. KEGG enrichment analysis of upregulated and downregulated DEGs in the NSPP plus radiation group compared to sham-irradiated control at 72 hours post-TBI (48 hours after NSPP application) **(f/g)**. Volcano plots of upregulated and downregulated DEGs in NSPP-treated groups, with or without radiation, compared to sham-irradiated control at 96 hours post-TBI **(h/i)**. KEGG enrichment analysis of upregulated and downregulated DEGs in the NSPP plus radiation group compared to sham-irradiated control at 96 hours post-TBI (72 hours after NSPP application) **(j/k)**. Volcano plot comparing upregulated and downregulated DEGs in the NSPP plus radiation group versus radiation-only group at 96 hour post-TBI **(l)**. KEGG enrichment analysis of upregulated and downregulated DEGs in the NSPP plus radiation group compared to irradiated control at 96 hours post-TBI (72 hours after NSPP application) **(m/n)**.

96 hours post-TBI is a crucial time point for observing GI-ARS recovery, characterized by the appearance of regenerative crypts and notable changes in gene expression and cellular pathways during this period [22]. In our study, the application of NSPP resulted in 101 significantly upregulated and 66 downregulated genes compared to sham-irradiated controls **(Figure 5h)**. Additionally, applying NSPP 24 hours post-TBI revealed a distinct gene signature, with 2616 upregulated and 1422 downregulated genes relative to the sham-irradiated group **(Figure 5i)**. KEGG enrichment analysis showed that the upregulated genes were involved in ECM-receptor interaction, focal adhesion, cytokine-cytokine receptor interaction, complement and coagulation cascade, and protein digestion and absorption **(Figure 5j)**. In contrast, the downregulated DEGs were associated with metabolism **(Figure 5k)**. Furthermore, NSPP application after TBI led to 24 significantly upregulated and 8 downregulated genes compared to the radiation-only group **(Figure 5l)**. The upregulated genes were linked to neuroactive ligand-receptor interaction and protein digestion and absorption **(Figure 5m)**, while the downregulated DEGs were associated with metabolism and IL-17 signaling pathway **(Figure 5n)**.

### Lgr5+ Intestinal Stem Cell Gene Signature changes during recovery following NSPP mitigation

Lastly, in order to investigate the impact of NSPP mitigation on ISC recovery, we compared the DEGs at 72 and 96 hours after TBI compared to their corresponding sham-irradiated controls with the list of *Lgr5^+^* ISC signature genes. Our analysis revealed a progressive and robust activation of these ISC signature genes, with the number of activated genes increasing as the recovery process advanced following NSPP treatment **(Figure 6)**. Importantly, this activation became more pronounced over time when compared to the effects of radiation alone **(Figure 3)**. These findings suggest that NSPP treatment significantly enhances the recovery of the *Lgr5+* ISC population, which is critical for effective tissue regeneration and repair. Among the key findings, *Psrc1* expression was notably higher following NSPP treatment, emphasizing its potential role in mediating the observed regenerative effects.

**Figure 6.**
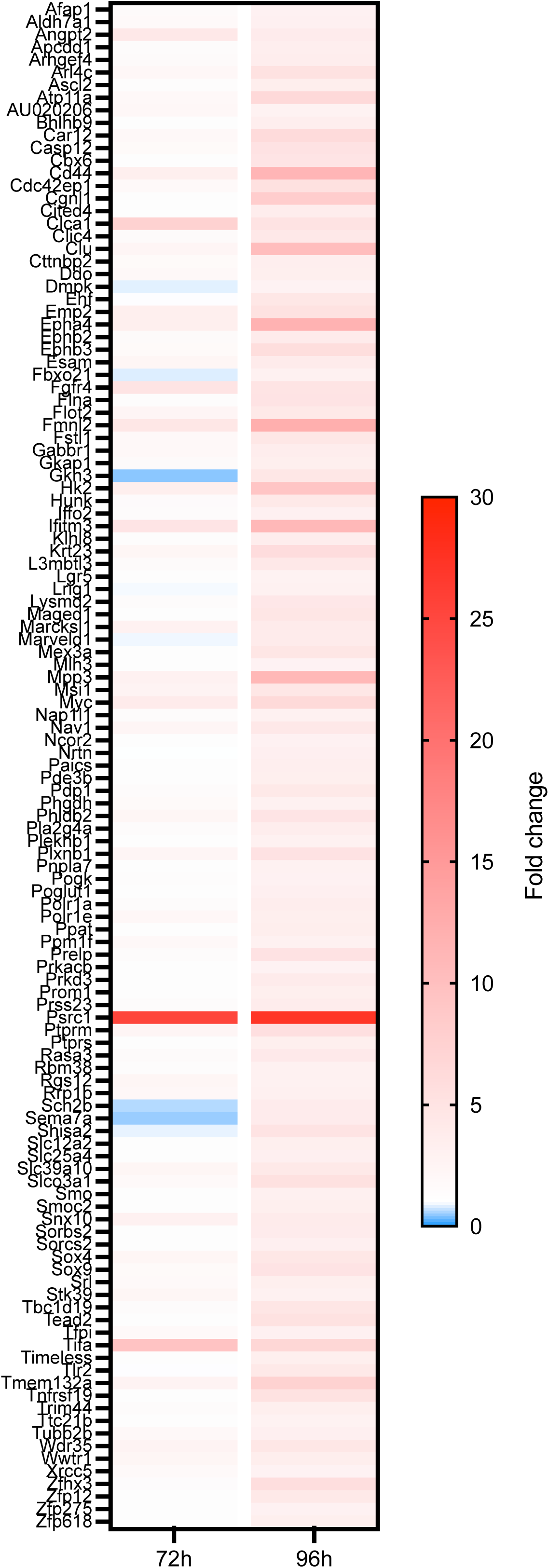
Changes in *Lgr5+* Intestinal Stem Cell Gene Signature during recovery following NSPP mitigation. Heatmap illustrating the enrichment of *Lgr5+* intestinal stem cell-associated genes among the up-regulated differentially expressed genes in the NSPP plus radiation group at 72 and 96 hours post-TBI compared to the corresponding sham-irradiated controls.

Further analysis using gene signatures associated with isthmus progenitor cells revealed that the intestinal stemness-associated gene *Ly6a* was already upregulated at 28 hours after TBI (4 hours after NSPP application), marking an early regenerative response. At 72 hours after TBI (48 hours after NSPP application), we observed a significant upregulation of *Areg* and *Anxa2*. At 96 hours after TBI (72 hours after NSPP application), we found further increases in the expression of *Ncl, S100a6,* and *Clu*. These findings underscore the ability of NSPP to increase the expression of key ISC-associated genes, highlighting its potential to enhance tissue regeneration and repair in the irradiated small intestines. The progressive upregulation of these critical genes provides a molecular basis for the observed improvements in recovery and reinforces the therapeutic potential of NSPP in mitigating radiation-induced damage.

## Discussion

In this study, we provide a comprehensive characterization of the transcriptomic changes in the murine duodenum following exposure to a lethal dose of total body irradiation. Our findings reveal dynamic shifts in gene expression over time, highlight the complex and coordinated response of intestinal tissues to radiation-induced damage, and underscore the therapeutic potential of NSPP in mitigating GI-ARS.

A key observation in the study was the activation of ISC-associated genes, such as *Lgr5+* population signatures, which were well-known to play a crucial role in epithelial repair and regeneration. However, *Lgr5+* ISCs are known to be highly sensitive to radiation and other external stimuli [23, 24], often leading to their depletion following exposure. This raises important questions about the involvement of alternative cell populations in maintaining and restoring intestinal integrity under severe damage conditions. One recent study has highlighted Tuft cells as a potential alternative regenerative population after radiation exposure. Tuft cells, which were traditionally considered as long-lived, postmitotic cells, exhibited an unexpected ability to revert to a stem-like state following radiation exposure [25]. These cells are highly resistant to radiation-induced damage and capable to generate organoids containing all intestinal epithelial cell types. Similarly, a recently study described that the *Fgfbp1+* ISC population in the upper crypt region generates progeny, propagating bi-directionally along the crypt-villus axis and replenishes *Lgr5+* cells in the crypt base. The high-resolution single cell profiling and *in vivo* lineage tracing in this report demonstrated that isthmus cells possess functional stem cell properties and participate in cellular turnover during both homeostasis and regeneration. Additional evidence of plasticity comes from *Dll1+* crypt cells and quiescent +4 label-retaining cells (LRCs) [26–28]. Under homeostatic conditions, *Dll1+* cells generate short-lived secretory clones, while quiescent +4 LRCs co-express *Lgr5* and markers of secretory lineages, such as Paneth and enteroendocrine cells. Both populations have been shown to regain stemness and contribute to regeneration following injury. These findings illustrate the hierarchical and flexible interplay of various cell types during tissue repair, adding new dimensions to our understanding of intestinal regeneration.

Our transcriptomic analysis revealed the activation of gene signatures associated with traditional *Lgr5+* ISC populations, suggesting that *Lgr5+* cells contribute to recovery and regeneration following radiation exposure. Additionally, the upregulation of the *Areg* and *Clu* genes – markers of isthmus progenitor cells – supports the presence of a reserve stem cell population likely involved in recovery and regeneration. More importantly, NSPP treatment further amplified the elevation of *Lgr5^+^* gene signatures and those associated with isthmus progenitor cells as early as 28 hours after TBI (4 hours after NSPP application). Furthermore, the expression of stemness-related genes such as *Ly6a, Areg, Anxa2, Ncl, S100a6,* and *Clu* was increased after NSPP treatment, indicating that NSPP enhances stem cell activation and the regenerative capacity of the intestinal epithelium following radiation-induced injury.

Among the genes upregulated during recovery in our study, *Psrc1* emerged as a particularly noteworthy candidate for its potential role in intestinal regeneration. *Psrc1* encodes a proline-rich protein regulated by the tumor suppressor protein p53 [29, 30]. It plays a crucial role in cell cycle progression and mitotic spindle function, both essential processes for ISC proliferation and differentiation. *Psrc1* marks a subset of ISCs enriched with MHC class II molecules involved in antigen presentation and interactions with immune cells [31]. These ISC-III cells, which are progressively more differentiated, also possess the capacity to de-differentiate and regain stem-like properties. In our study, NSPP treatment significantly amplified *Psrc1* expression, suggesting that this gene may serve as a key mediator of stem cell activation and epithelial repair. The increased expression of *Psrc1* likely contributes to the enhanced regenerative capacity observed with NSPP treatment, emphasizing its critical role in stem cell-mediated recovery. Future studies should explore the mechanistic role of *Psrc1* in ISC biology, particularly its influence on cellular plasticity and its interactions with other regenerative pathways and compartments within the gastrointestinal system. Understanding how *Psrc1* contributes to ISC functionality and its cross-talk with immune cells may reveal novel strategies for enhancing intestinal repair and mitigating damage following radiation or other injuries.

The parallels between radiation-induced plasticity in the intestine and in cancer biology are also noteworthy. Our previous research in glioblastoma [32, 33], head and neck cancer [34], and breast cancer [35, 36] demonstrated that radiation can induce transient multipotent states, leading to increased cellular plasticity. Similarly, the regenerative ability of intestinal cells, including Tuft cells [25] and other non-canonical populations [21, 22, 27], highlights the fundamental importance of plasticity as a response to radiation damage. While this plasticity facilitates tissue repair, it also raises concerns about potential risks, such as dysregulated repair or tumorigenesis, that warrant further investigation.

Looking forward, future research should validate these findings in other regions of the intestine to determine whether similar mechanisms of regeneration apply along the gastrointestinal tract. Additionally, assessing the long-term effects of NSPP treatment, including its impact on survival, tissue functionality, and potential side effects, will be crucial for evaluating its therapeutic potential. Investigating the interactions between canonical ISCs, alternative stem cell populations, and niche components such as immune and endothelial cells will also enhance our understanding of the complex intestinal response to radiation damage.

While our study provides valuable insights into the transcriptomic changes in the murine duodenum following TBI and highlights the potential of NSPP as a radiation mitigator, several limitations should be considered. These findings need to be validated in other regions of the intestine to determine whether similar regenerative mechanisms are consistent throughout the gastrointestinal tract. Additionally, assessing the long-term effects of NSPP treatment, including its impact on survival, tissue functionality, and potential side effects, is crucial for evaluating its therapeutic viability. Finally, the use of bulk RNA sequencing in this study does not provide information about the interactions between different cellular compartments within the epithelium, such as immune cells and endothelial cells. Single cell RNA sequencing or spatial transcriptomics could be beneficial as they have the potential to unravel complex cellular interactions by providing high-resolution data on gene expression at the individual cell level and preserving spatial context within the tissue. This could lead to a more comprehensive understanding of the cellular dynamics and molecular mechanisms involved in intestinal regeneration and how NSPP influences these processes following radiation exposure.

## Conclusions

In summary, our study highlights the complex and dynamic molecular mechanisms underlying intestinal recovery following radiation exposure. The contributions of *Lgr5^+^* intestinal stem cells, and possibly other plasticity-driven cell populations, are essential for tissue repair. NSPP’s ability to amplify these regenerative processes underscores its therapeutic potential. These findings enhance our understanding of radiation-induced damage and recovery, providing a foundation for developing targeted therapies to mitigate radiation injuries and improve outcomes in radiological emergencies.

## Acknowledgements

Not applicable.

## Authors’ contributions

FP conceived and supervised the overall project, designed the experiments. KB and LH performed the experiments, FP and LH analyzed the data and drafted the manuscript. All authors contributed to reviewing/editing the final version of the manuscript.

## Data Availability

Research data are stored in an institutional repository and will be shared upon request to the corresponding author.

